# Oral copper-methionine decreases matrix metalloproteinase-2 activity in the liver and brain of broiler chickens subjected to cold stress for ascites incidence

**DOI:** 10.1101/2023.10.14.562342

**Authors:** Mina Bagheri Varzaneh, Hamidreza Rahmani, Rahman Jahanian, Amir Hossein Mahdavi, Corinne Perreau, Stéphane Brézillon, François-Xavier Maquart

## Abstract

Copper plays an antioxidant role in biological reactions. This study examined the impact of copper-methionine supplementation on the matrix metalloproteinase-2 (MMP-2) activity and gene expression in the liver and brain of broiler chickens subjected to cold temperature. A total of 480 broiler chickens were assigned to 6 groups and reared under either low (15-19 ºC) or normal temperature (25-28ºC) and fed a basal diet enriched with different concentrations of copper-methionine (Cu-Met) supplementation (0, 100 or 200 mg.kg^-1^). Ascites was exclusively observed in broiler chickens kept in low temperature and fed with basal diet without Cu-Met during the seventh week, identified by the presence of abdominal fluid accumulation. Broilers’ livers and brains were separated for MMP-2 and tissue inhibitor of metalloproteinase-2 (TIMP-2) analysis. Results of gelatin zymography on these samples demonstrated that incidence of ascites was associated with increased MMP-2 levels in liver and brain. MMP-2 activity assay confirmed the results obtained by zymography. RT-qPCR experiments revealed an upregulation in the mRNA expression of MMP-2. In contrast, the treatments did not induce significant alterations in TIMP-2 levels. Results suggest that oral copper-methionine can decrease the ascites occurrence and might be useful for prevention of ascites in broiler chickens.

## Introduction

Matrix metalloproteinases (MMPs) are a large group of zinc-dependent enzymes that degrade extracellular matrix (ECM) proteins such as collagens and elastin. These are the triggers for activation the different signaling pathways [1,2]. ECM degradation is necessary to cell invasion, cell motility and proliferation in different diseases [3,4]. In this way, the role of MMPs, especially gelatinase-A (MMP-2) and gelatinase-B (MMP-9) as major agents in basement membrane degradation has been reported in the pathophysiology of fatty liver disease [5]. In addition, MMPs can stimulate the production and secretion of growth factors, such as vascular endothelial growth factor (VEGF) [6,7]. VEGF itself increases vascular permeability and plays a critical role in ascites formation [8,9]. Furthermore, VEGF is expressed in chronic liver diseases such as fatty liver disease and liver cirrhosis [5,10]. In addition to these diseases, the activation of MMPs and VEGF have some pathogenic effect on brain and may promote tumor formation in the central nervous system (CNS) [11,12]. Tissue inhibitors of metalloproteinases (TIMPs) are originally characterized as inhibitors and key regulators of MMPs, therefore TIMPs can minimize degradation of matrix. Degradation of ECM depends on the balance between TIMPs and MMPs activities [13].

Copper is an essential trace element as a cofactor in enzymes such as superoxide dismutase and lysyl oxidase. Due to its critical role in mitochondrial function and the peroxisomal beta-oxidation of fatty acids, copper deficiency leads to impaired mitochondrial function and increased oxidative stress. Copper deficiency disrupts the normal functioning of mitochondria, resulting in compromised energy production and an imbalance in oxidative stress levels [14]. Lysyl oxidase is a cuproenzyme that plays a critical role for crosslinking of the extracellular matrix proteins such as elastin and collagen. Superoxide dismutase is a metalloenzyme that plays a major role in protection against oxidative stress by degrading reactive oxygen species [15,16]. It has been demonstrated that the level and form of copper changed the liver performance and copper profile in broiler chickens [17]. Despite much information about the role of copper in biological reactions, no study was reported about the effects of copper-methionine on the liver and brain in broiler chickens. In this way, we investigated the effect of Cu-Met supplementation on TIMP-2 and gelatinases expression and activation in liver and brain of ascitic broiler chickens. Our findings can be potentially helpful for ascites prevention.

## Materials and Methods

### Experimental design

Four hundred eighty commercial broilers (*Ross 308*) were divided into six treatment groups. Treatments included two growing temperatures: either standard (25-28 ºC) or low (15-19 ºC). Broilers in each temperature condition were fed with a basal diet with three added copper levels of Cu-Met complex (0, 100 and 200 mg.kg^-1^ of diet). At seventh week, the livers, and brains of four chickens from each group were collected and frozen at -80 ºC.

### Gelatin and reverse zymographies

ProMMP-2 and TIMP-2 were assessed by gelatin substrate zymography and gelatin reverse zymography, respectively. Gelatin zymography was performed according to Kleiner and Stetler-Stevenson [18], slightly modified. At first, homogenized samples were centrifuged at 4ºC for 10 minutes. Protein quantification kit (Bradford assay) were used to calculate the protein concentration in supernatants. Samples (20 μg of proteins for each sample) were loaded on SDS-PAGE (8% polyacrylamide minigel with 0.1% gelatin). After electrophoresis, the gel was washed in Triton X-100 and incubated overnight at 37°C. Then, the gel was stained with a solution of Coomassie blue. Destaining was performed in 10% acetic acid and 20% methanol to detect the MMP-2 and MMP-9 bands, which appear as clear bands on a blue background [18].

For reverse zymography, samples were separated by 15% SDS-PAGE supplemented with proMMP-2 and 0.1% gelatin. In this method, TIMP-2 prevented the local degradation of gelatin by proMMP-2 during incubation, so that dark bands in a white background demonstrated TIMP-2 activity area.

Conditioned medium from human fibrosarcoma HT-1080 cell line was run as a positive control for MMP-2, MMP-9 and TIMP-2 detection in gelatin zymography and gelatin reverse zymography. The bands were quantified by ChemiDoc MP (BioRad) [19,20].

### Gene expression and enzyme activity assay

After RNA extraction, the quality and concentration of RNA were measured by spectrophotometric analysis (NanoVue Plus, USA). RNA was reverse transcribed to cDNA and PCR was performed for 35 cycles. During electrophoresis PCR products were separated by 2% agarose gel and ethidium bromide was used for staining the gel. β-actin mRNA expression was used for normalization. Reactions were carried out in duplicate with identical concentrations of cDNA. MMP-2 mRNA expression was analyzed by the comparative CT method [20].

Analyses of MMP-2 activity were performed by a fluorescence assay kit (SensoLyte® 520 MMP-2 kit, AnaSpec, San Jose, USA). The fluorescence of the substrate cleavage product was determined at excitation/emission = 490 nm/520 nm with a Perkin-Elmer spectrofluorometer [21,22].

### Statistical Analysis

All data were analyzed using SAS software (1998). Statistical significance was distinguished by the General Linear Model (GLM) procedure. The experiment was performed in 2 × 3 factorial with 2 levels of temperature and 3 levels of Cu-Met and treatments were assigned at completely randomized block design. The data was presented as the mean ± S.E and p< 0.05 declared as statistically significant.

## Results

### MMP-2 activity and proMMP-2 levels in liver and brain

Results of gelatin zymography showed that only proMMP-2 was present in the liver and brain extracts, without any detectable proMMP-9.

Clinical ascites happened in broilers fed with the basal diet (without Cu-Met supplementation) in cold conditions at seventh week. ProMMP-2 levels and MMP-2 activity in the liver extract were significantly increased in ascitic broiler chickens. Cu-Met supplementation (100 and 200 mg.kg^-1^) significantly decreased proMMP-2 levels and MMP-2 activity in cold temperature treatments whereas no significant alteration was observed in normal temperature (Figure 1,2).

**Figure 1.**
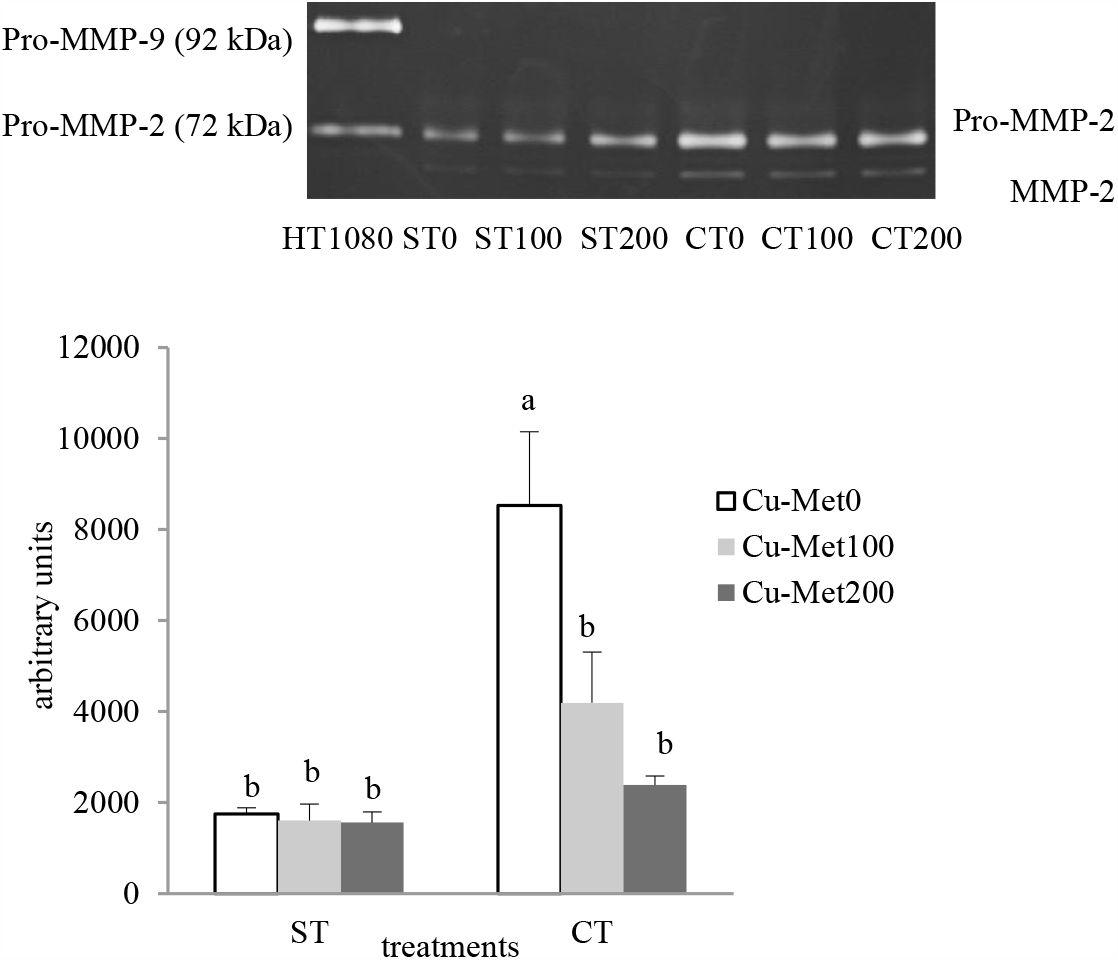
Effects of treatments on proMMP-2 in the broilers’ liver at seventh week. a) proMMP-2 bands were detected on gelatin zymogram. Treatments included: standard temperature, Cu-Met 0 mg.kg^-1^ (ST0); standard temperature, Cu-Met 100 mg.kg^-1^ (ST100); standard temperature, Cu-Met 200 mg.kg^-1^ (ST200); cold temperature, Cu-Met 0 mg.kg^-1^ (CT0); cold temperature, Cu-Met 100 mg.kg^-1^ (CT100); cold temperature, Cu-Met 200 mg.kg^-1^ (CT200). b) Quantitative analysis of gelatin zymogram of proMMP-2. ^a,b^ Means with different letters are significantly different (P< 0.05).

**Figure 2.**
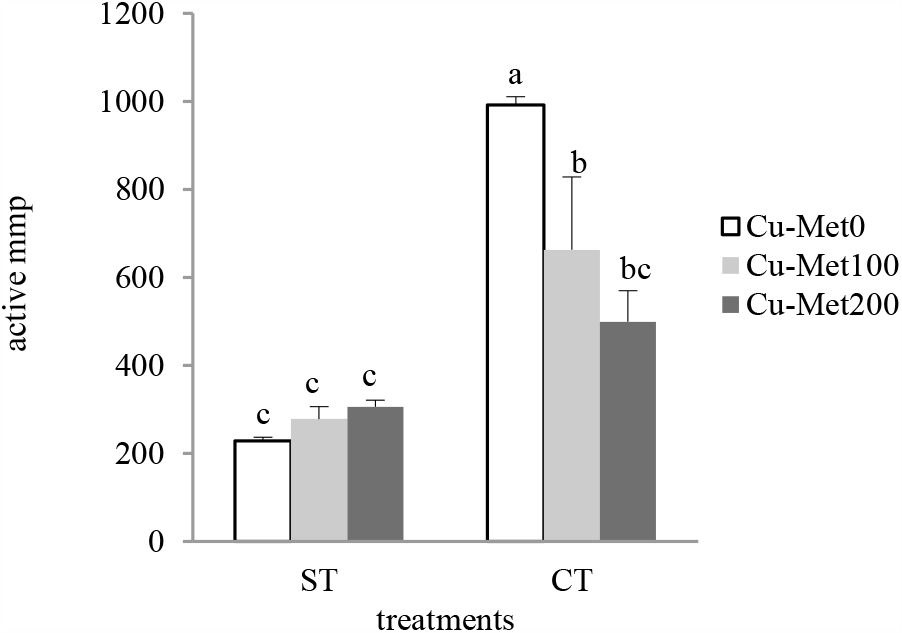
MMP-2 activity fluorometric assay showing the effects of Cu-Met and cold temperature in liver extracts of broilers at seventh week. Treatments included: standard temperature, Cu-Met 0 mg.kg^-1^ (ST0); standard temperature, Cu-Met 100 mg.kg^-1^ (ST100); standard temperature, Cu-Met 200 mg.kg^-1^ (ST200); cold temperature, Cu-Met 0 mg.kg^-1^ (CT0); cold temperature, Cu-Met 100 mg.kg^-1^ (CT100); cold temperature, Cu-Met 200 mg.kg^-1^ (CT200). ^a,b,c^ Means with different letters are significantly different (P < 0.05).

In the brain of broiler chicken grown under cold temperature at seventh week, proMMP-2 levels were increased. The brain of chickens reared in normal conditions showed the lowest proMMP-2 levels. Interestingly, feeding with copper-methionine supplementation decreased proMMP-2 to levels similar to that observed in the brain of chickens grown under normal temperature treatments (Figure 3). This was confirmed by fluorometric assay of MMP-2 activity (Figure 4).

**Figure 3.**
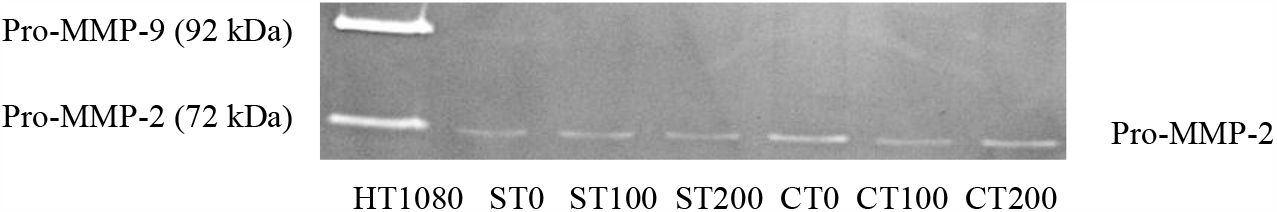
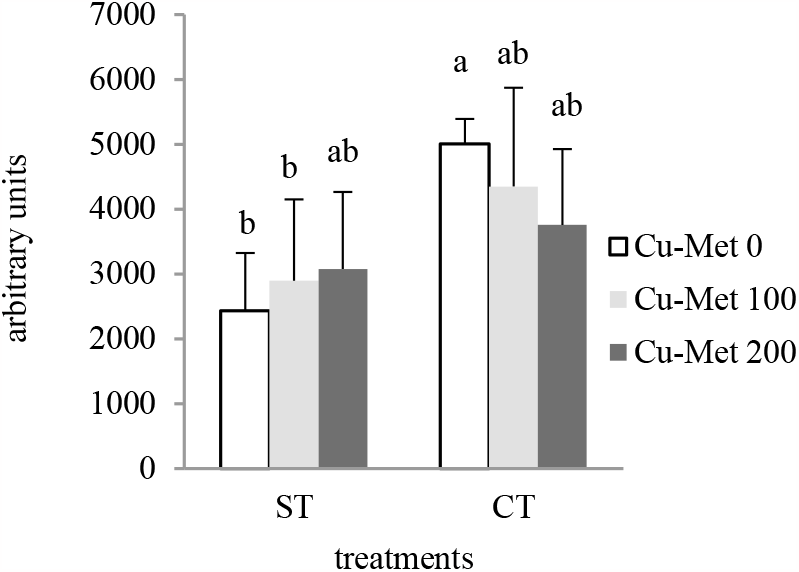
Effects of treatments on proMMP-2 in the broilers’ brain at seventh week: a) proMMP-2 bands were detected on gelatin zymogram. Treatments included: standard temperature, Cu-Met 0 mg.kg^-1^ (ST0); Standard temperature, Cu-Met 100 mg.kg^-1^ (ST100); Standard temperature, Cu-Met 200 mg.kg^-1^ (ST200); Cold temperature, Cu-Met 0 mg.kg^-1^ (CT0); Cold temperature, Cu-Met 100 mg.kg^-1^ (CT100); Cold temperature, Cu-Met 200 mg.kg^-1^ (CT200). b) Quantitative analysis of gelatin zymogram of proMMP-2. ^a,b^ Means with different letters are significantly different (P< 0.05).

**Figure 4.**
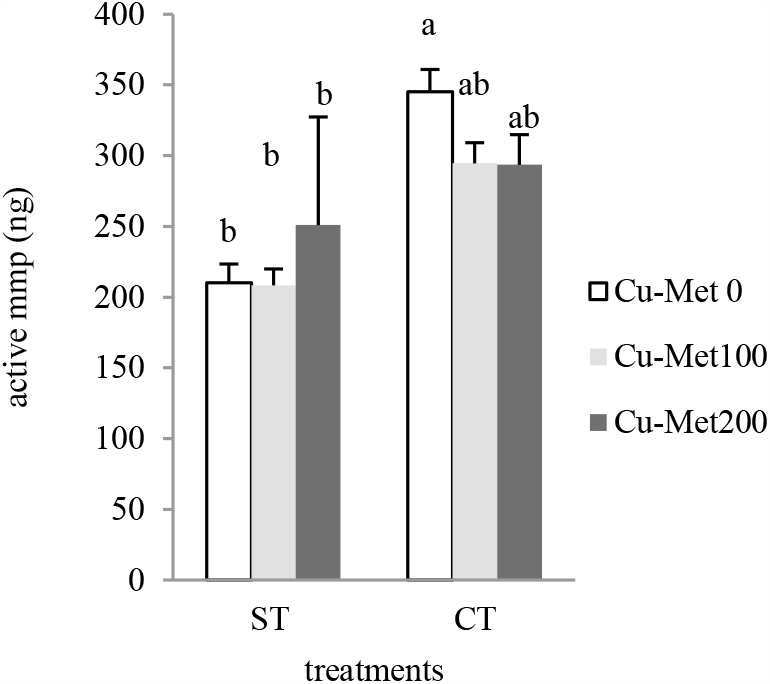
MMP-2 activity fluorometric assay showing the effects of Cu-Met and cold temperature in brain extracts of broilers at seventh week. Treatments included: standard temperature, Cu-Met 0 mg.kg^-1^ (ST0); Standard temperature, Cu-Met 100 mg.kg^-1^ (ST100); Standard temperature, Cu-Met 200 mg.kg^-1^ (ST200); Cold temperature, Cu-Met 0 mg.kg^-1^ (CT0); Cold temperature, Cu-Met 100 mg.kg^-1^ (CT100); Cold temperature, Cu-Met 200 mg.kg^-1^ (CT200). ^a,b^ Means with different letters are significantly different (P < 0.05).

### MMP-2 mRNA expression in liver and brain

MMP-2 mRNA expression was also done using standard RT-PCR and real time PCR analysis. In the liver of broilers experiencing the ascites, RTq-PCR showed higher level of MMP-2 gene expression. Feeding supplementation with Cu-Met decreased MMP-2 mRNA expression in the liver of broiler chicken reared under cold stress (Figure 5).

**Figure 5.**
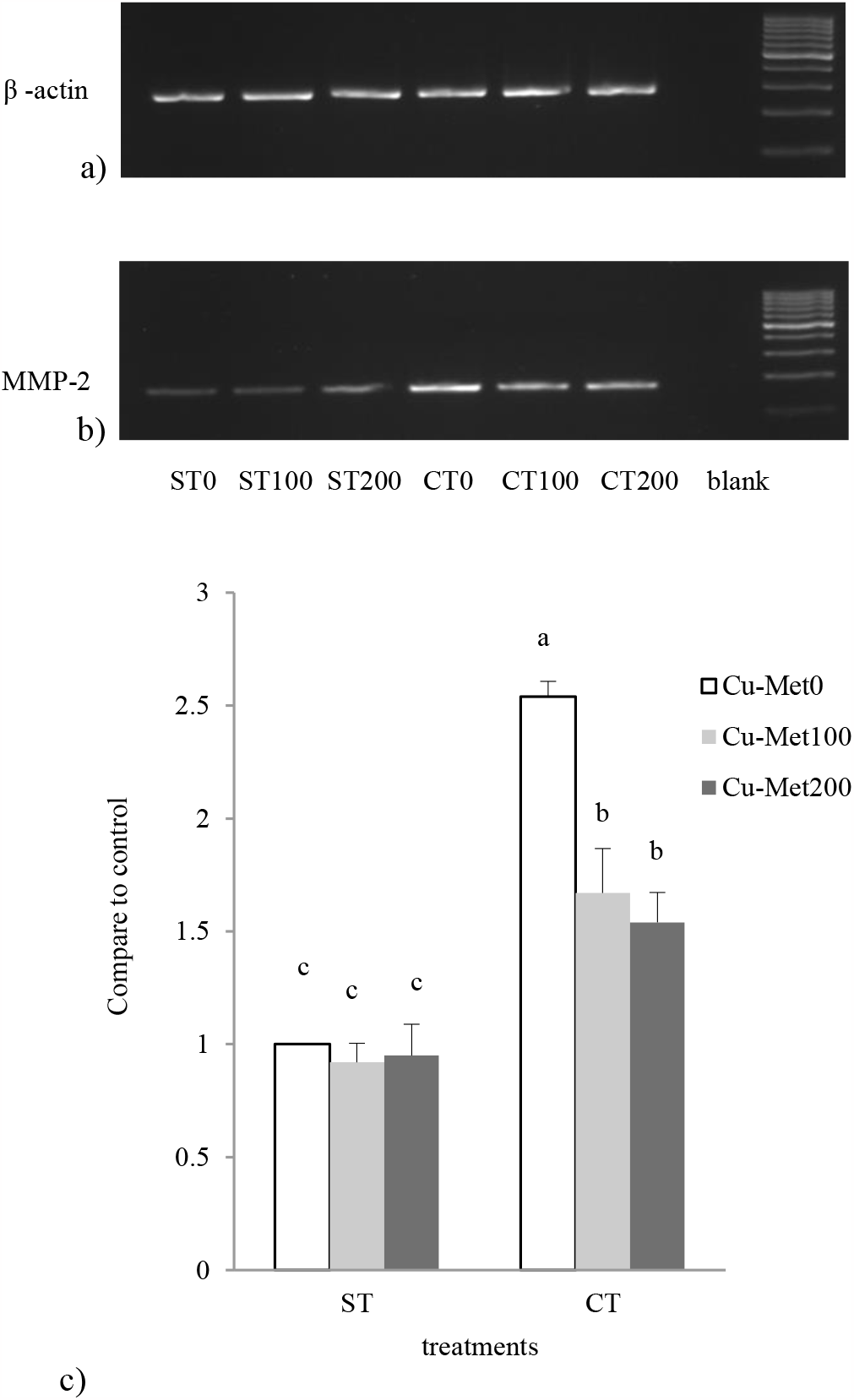
(a) Standard RT-PCR for β -actin gene expression in the liver of broiler chicken at seventh week. (b) Standard RT-PCR for MMP-2 gene expression. Treatments included: standard temperature, Cu-Met 0 mg.kg^-1^ (ST0); standard temperature, Cu-Met 100 mg.kg^-1^ (ST100); standard temperature, Cu-Met 200 mg.kg^-1^ (ST200); cold temperature, Cu-Met 0 mg.kg^-1^ (CT0); cold temperature, Cu-Met 100 mg.kg^-1^ (CT100); cold temperature, Cu-Met 200 mg.kg^-1^ (CT200). (c) Real-time PCR analysis in broilers’ liver at seventh week. ^a,b,c^ Means with different letters are significantly different (P < 0.05).

In the brain of broilers kept in the normal conditions, MMP-2 mRNA did not change at any copper concentration. On the contrary, in the brain of broilers reared in cold conditions, MMP-2 mRNA expression was increased but it decreased with Cu-Met diet supplementation 100 and 200 mg.kg^-1^ (Figure 6).

**Figure 6.**
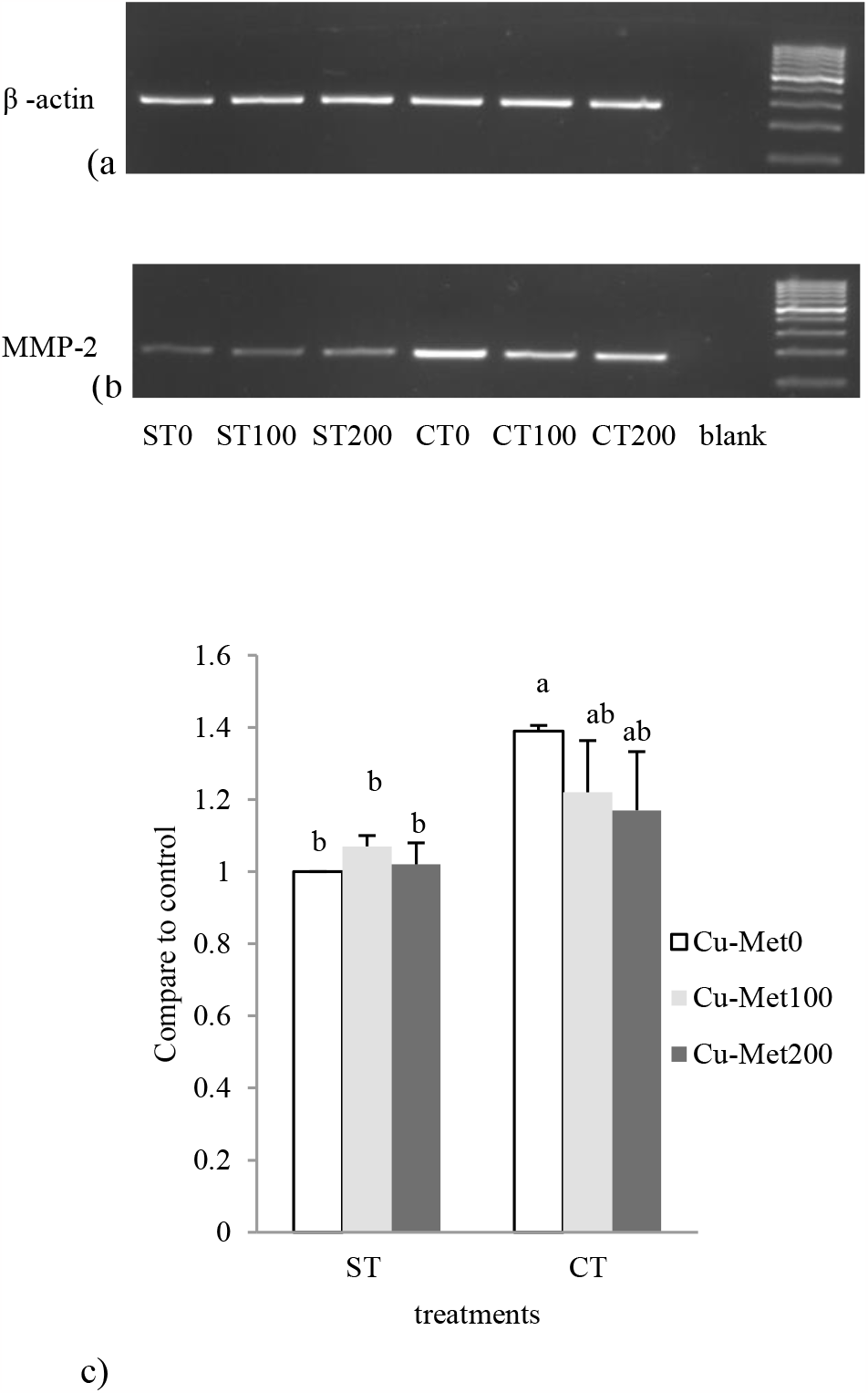
(a) Gene expression of β -actin in the brain analyzed by standard RT-PCR. (b) Gene expression of MMP-2 analyzed by standard RT-PCR at seventh week. Treatments included: standard temperature, Cu-Met 0 mg.kg^-1^ (ST0); Standard temperature, Cu-Met 100 mg.kg^-1^ (ST100); Standard temperature, Cu-Met 200 mg.kg^-1^ (ST200); Cold temperature, Cu-Met 0 mg.kg^-1^ (CT0); Cold temperature, Cu-Met 100 mg.kg^-1^ (CT100); Cold temperature, Cu-Met 200 mg.kg^-1^ (CT200). Blanks were prepared using a similar process without cDNA. (c) Relative quantification of MMP-2 mRNA expression by real-time PCR in broilers’ brain at seventh week. ^a,b^ Means with no common superscript letters are significantly different (p < 0.05).

### Tissue inhibitors of metalloproteinases levels in liver and brain

Only TIMP-2 was detected in the liver and brain extracts of all groups. TIMP-2 levels were evaluated using reverse gelatin zymography at seventh weeks. Results showed that low temperature and oral Cu-Met at different levels have no effect on TIMP-2 expression in liver (Figure 7). Levels of TIMP-2 in the brain of chickens reared under normal and cold conditions were not affected by Cu-Met supplementation (100 and 200 mg.kg^-1^) (Figure 8).

**Figure 7.**
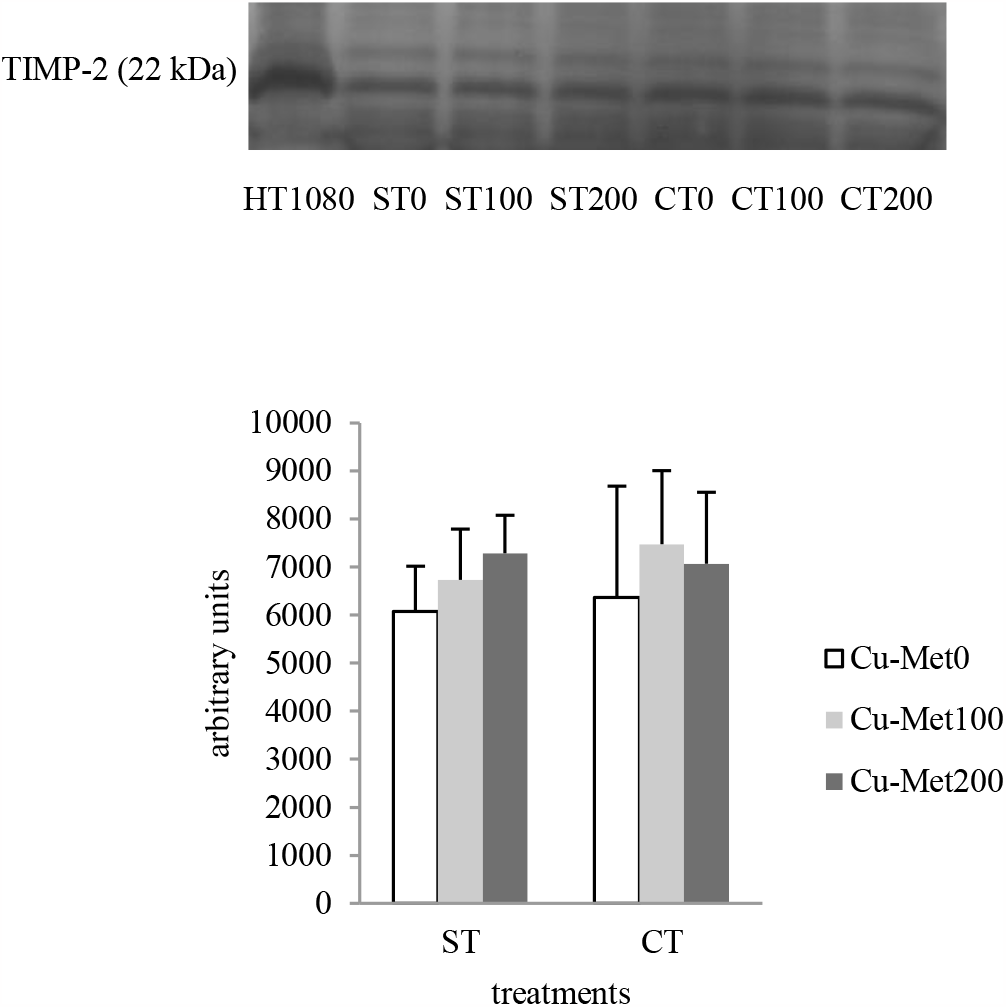
a) Effects of Cu-Met at different levels and cold temperature on TIMP-2 level in the broilers’ liver at seventh week. Treatments included: standard temperature, Cu-Met 0 mg.kg^-1^ (ST0); standard temperature, Cu-Met 100 mg.kg^-1^ (ST100); standard temperature, Cu-Met 200 mg.kg^-1^ (ST200); cold temperature, Cu-Met 0 mg.kg^-1^ (CT0); cold temperature, Cu-Met 100 mg.kg^-1^ (CT100); cold temperature, Cu-Met 200 mg.kg^-1^ (CT200). b) Quantitative analysis for TIMP-2 detection. No significant difference could be detected.

**Figure 8.**
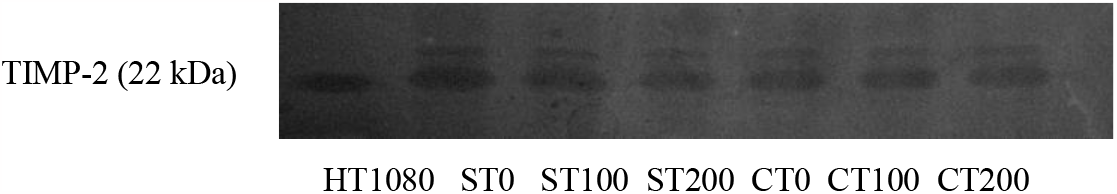
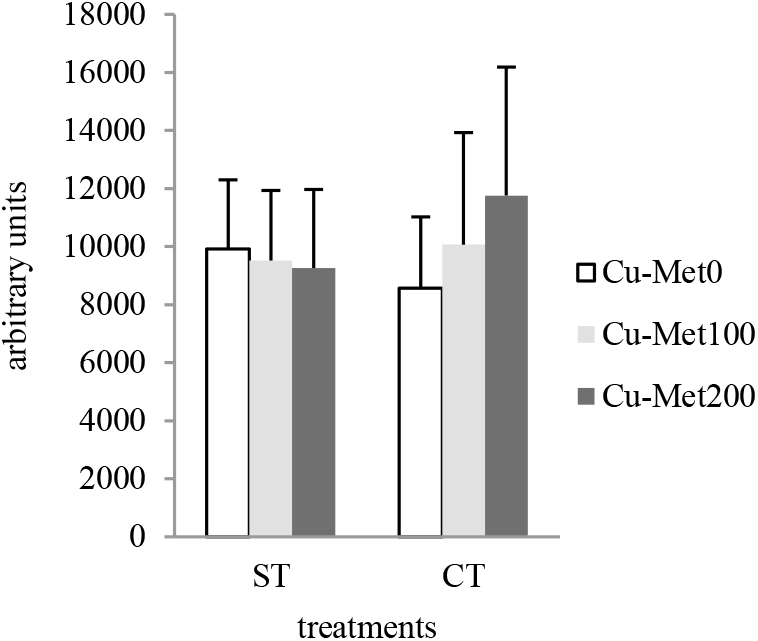
a) Effects of Cu-Met at different levels and cold temperature on TIMP-2 level in the broilers’ brain at seventh week. Treatments included: Standard temperature, Cu-Met 0 mg.kg^-1^ (ST0); Standard temperature, Cu-Met 100 mg.kg^-1^ (ST100); Standard temperature, Cu-Met 200 mg.kg^-1^ (ST200); Cold temperature, Cu-Met 0 mg.kg^-1^ (CT0); Cold temperature, Cu-Met 100 mg.kg^-1^ (CT100); Cold temperature, Cu-Met 200 mg.kg^-1^ (CT200). b) Quantitative analysis for TIMP-2 detection. No significant difference could be detected.

## Discussion

Today’s very fast-growing broilers have high metabolic rates imposing physiologically a high demand of oxygen which their cardiopulmonary system is unable to supply. Growing under cold temperature even exacerbates the growing conditions because it has devastating effects on mitochondrial processes. Indeed, oxygen deficiency in mitochondrion increases toxic compounds such as ROS in cells. As a consequence of oxidative stress, ascites can be induced by cold temperatures. It is a major problem in broilers and can account for up to 5% mortality in broiler flocks and decrease the performance, meat quality and overall production efficiency. In this way, ascites can pose a significant economic burden in broiler production [23,24].

Interactions between genetic parameters and environmental risk factors are major agents in ascites development. Scientific evidence suggests that ROS produced through lipid peroxidation plays an important role in ascites formation. Dietary antioxidants can exert a protective effect against ascites progression by either scavenging ROS or inhibiting cellular signaling enzymes that induce cell invasion and proliferation [4,25]. There is increasing evidence that MMPs, especially gelatinases, are involved in different diseases such as CNS cancers, liver diseases and ascites progression [12,26,27]. Increased activity of MMP-2 and MMP-9 has been reported in cell invasion and cancer formation [28]. According to Ozyigit *et al*. and Belotti *et al*., increased gelatinases activity could be implicated in the development of ascites. Ascites, which may result from high activity of gelatinases [7,29], is sometimes viewed as a symptom in CNS cancer and liver diseases [5,28]. Chemical MMP inhibitors such as tetracyclines have been investigated for their potential in treating neurological disease. However, the safety problems and the adverse effects of these components are typically observed over the medium and long term uses in clinical trials [30]. Therefore, the importance of antioxidant compounds in inhibiting MMPs becomes apparent. Lu et al. has reported that inhibiting reactive oxygen species (ROS) by rosmarinic acid decreased MMP-2 expression in hepatic stellate cells [31]. The catechins found in green tea have been shown to effectively inhibit MMP-2 activities in the glioblastomas cells [32]. Furthermore, there are some studies which have demonstrated that trace elements modulated MMPs activity and improved cell migration and invasion in different kinds of disease [33-35]. In this study, cold stress in broiler chickens induced ascites which was associated with increased MMP-2 activity in liver and brain, while copper-methionine supplementation alleviated the problem. Copper plays a crucial role in the functioning of essential antioxidant enzymes, such as superoxide dismutase (SOD), ceruloplasmin, cytochrome c oxidase (COX) and hephaestin [14]. It has been shown that copper, as a cofactor of superoxide dismutase (SOD), is involved in antioxidant reactions by preventing the destructive effects of ROS in liver and brain cells [36,37]. Copper deficiency has been found to be associated with dyslipidemia, including obesity and non-alcoholic fatty liver disease (NAFLD) [38,39]. Aigner *et al* has shown that the patients with NAFLD have lower concentration of copper in liver [40].

Wu et al. demonstrated that 30 mg.kg^-1^ Cu-Met improved feed conversion rate (FCR), reduced serum triglycerides and cholesterol levels and increased fat digestibility. Furthermore, adding Cu-Met to the diet has been found to increase the activities of glutathione peroxidase, ceruloplasmin, superoxide dismutase and immune indexes such as IgA and IgM in broiler chickens [41]. It is important to note that improper supplementation of copper can have negative consequences on liver health. Overdosing on copper can disrupt mitochondrial metabolism, as evidenced by the emergence of a recently discovered phenomenon known as cuproptosis. Cuproptosis refers to a type of regulated cell death that is dependent on copper levels in the cell. Therefore, maintaining an appropriate balance of copper is crucial to avoid detrimental effects on liver function and mitochondrial metabolism [42,43]. Deo et al. has reported that 100 mg.kg^-1^ Cu-Met improved the growth performance and cell-mediated immune response in broiler Japanese quails [44]. The organic source of copper has a higher bioavailability and demonstrated superior effectiveness in promoting growth and decrease the cholesterol levels compared to other copper sources in broiler chickens [45].

Current results on decreased MMP-2 activity and gene expression in broilers liver and brain after copper supplementation are in line with our previous work reporting that oral copper-methionine reduced proMMP-2 levels and MMP-2 activity in lungs and heart of broiler chickens [21,22]. Copper is particularly important for the development of the central nervous system and plays an important role in neurotransmitter biosynthesis and proper neurological functioning. Insufficient copper levels can lead to incomplete neurological development, while excessive copper can generate free radicals through the Haber-Weiss reaction, which induce damage to neurons [46]. Furthermore, recent studies demonstrated that copper enhances the therapeutic effects of drugs on central nervous system tumor and liver diseases [25,47-49].

TIMPs can bind MMPs and block their enzymatic function [50]. Daniele et al. has been reported the involvement of MMP-2, MMP-9, TIMP-1, and TIMP-2 in degrading the extracellular matrix in hepatocellular carcinoma (HCC). Additionally, in chronic liver diseases such as cirrhosis and hepatitis B and C, the expression of MMP-9, MMP-2, TIMP-1, TIMP-2 and MMP-2,9/TIMP-1,2 ratio can serve as a useful parameter for monitoring disease progression and predicting the development of diseases [51]. In this study, TIMP-2 activity was not different between the different groups. These data show that MMP-2 activity changes were independent of TIMP-2 but depended mainly on MMP-2 gene expression and MMP-2 activation.

To our knowledge, no previous study has investigated the effect of Cu-Met as an organic source of copper in liver and brain in ascitic broilers and its effects on MMP-2 expression and activity. Our results indicate that a Cu-Met diet supplementation of 100 or 200 mg.kg^-1^ inhibits MMP-2 gene expression and activity. This inhibiting effect could be important for controlling the progression of ascites. Diet supplementation with 100 and 200 mg.kg^-1^ of Cu-Met did not show any toxic effects in broilers and the results between two levels of Cu-Met supplementation were the same.

## Conclusion

Cold stress increased the incidence of ascites in broiler chickens, which was associated with increased MMP-2 gene expression and activity in the liver and brain of affected birds. Interestingly, supplementation of 100 and 200 mg.kg^-1^ Cu-Met reduced MMP-2 gene expression and MMP-2 activity. Results suggest performing more detailed studies to elucidate the molecular mechanisms and signaling pathways involved in positive effects of high levels of copper supplementation. Totally, we suggest that copper-methionine diet supplementation could be tested in ascites prevention in broiler chickens.

## Compliance with Ethical Standards

### Author Contributions

Writing—original draft preparation; M.BV. Writing—review and editing; F.X.M., S.B., C.P., H.R., A.M., and R.J.; Supervision, F.X.M. and H.R. All authors have read and agreed to the published version of the manuscript.

### Funding

This research received no external funding

### Institutional Review Board Statement

The study was conducted according to the Guide for the Care and Use of Laboratory Animals published by the National Institutes of Health (NIH Publications No. 8023, revised 1978).

### Informed Consent Statement

Not applicable

### Data Availability Statement

The data presented in this study are available on request from the corresponding author.

